# Kernel integration by Graphical LASSO

**DOI:** 10.1101/2020.03.11.986968

**Authors:** Sarah Kristine Nørgaard, Kristoffer Linder-Steinlein, Anders Ulrik Eliasen, Jakob Stokholm, Bo L. Chawez, Klaus Bønnelykke, Hans Bisgaard, Age K. Smilde, Morten A. Rasmussen

## Abstract

Integration of unstructured and very diverse data is often required for a deeper understanding of complex biological systems. In order to uncover communalities between heterogeneous data, the data is often harmonized by constructing a kernel and numerical integration is performed. In this study we propose a method for data integration in the framework of an undirected graphical model, where the nodes represent individual data sources of varying nature in terms of complexity and underlying distribution, and where the edges represent the partial correlation between two blocks of data. We propose a modified GLASSO for estimation of the graph, with a combination of cross-validation and extended Bayes Information Criterion for sparsity tuning. Furthermore, hierarchical clustering on the weighted consensus kernels from a fixed network is used to partitioning the samples into different classes. Simulations show increasing ability to uncover true edges with increasing sample size and *signal to noise*. Likewise, identification of non existing edges towards disconnected nodes is feasible. The framework is demonstrated for integration of longitudinal symptom burden data from the 2nd and 3rd year of life with 21 diseases precursors as well as the development of asthma and eczema at the age of 6 years from 403 children from the COPSAC2010 mother-child cohort, suggesting that maternal predisposition as well as being born preterm indirectly lead to higher risk of asthma via increased respiratory symptom burden.

## 1 | INTRODUCTION

Understanding complex biological systems often requires the need to integrate data of very diverse nature, such as data with variable dimensionality and distribution, with a mixture of longitudinal or cross sectional features. Given such a set of distributed sources of information pertaining to the same set of samples is often tackled by harmonizing the data heterogeneity by construction of so-called kernels followed by a numerical integration aiming at uncovering commonality [1]. Kernel transformation resembles the concept of characterizing some information entity 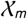 from *n* individuals into a symmetric kernel 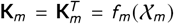, where 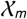 can be anything from univariate random variables, multivariate, tables of counts, time-to-event, etc. [2, 3]. After kernel-harmonization 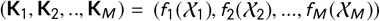, these matrices can be integrated by factorization such as INDSCAL [4] to reveal common structure between the *M* data layers, as well as uncovering the multivariate distribution of the *n* samples. While such factorization models are powerful in estimating patterns of correlation, they do not infer the structural topology related to the conditional dependency between layers of data. Aben *et al* 2018 developed a framework for inferring the conditional dependency between data layers based on the kernel-to-kernel (matrix) correlations, referred to as the RV coefficient [5, 6], and used this for structure revealing from seven data layers covering genotype to phenotype information from a cancer study [1].

In this study we propose a method for data integration in the framework of an undirected graphical model. Let 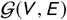 be a representation of a graph, where *V* and *E* are the vertices (nodes) and edges between *M* sources of information. In particular for this study, the nodes represent individual data sources of varying nature in terms of complexity and underlying distribution, and the edges represent the partial correlation between two blocks of data (conditioning on all other data blocks). We propose a modified graphical lasso for estimation of the graph, with a combination of cross-validation and extended Bayes Information Criterion [7] for sparsity tuning. Naturally, the choice of functions *f_m_* for kernel transformation, inherently affects the upstream results, and depending on the nature of the data there is a vast source of methods for such transformations (see e.g. [8] for details). For this purpose, we utilize data from a clinical childhood cohort with information on recurrent periods of specific diagnoses and disease remission (amoung other sources), we propose kernel transformations for such data. Having established a graph between the layers of data, we propose a clustering method based on weighted consensus kernels and hierachical clustering for partitioning of the samples into different classes. The method is demonstrated on simulated as well as real data pertaining to symptom burden of common childhood diseases across the 2^nd^ and 3^rd^ year of life in relation to exposures as well as the development of asthma at age 6 years from the COPSAC2010 mother-child cohort [9, 8].

The paper is organized with a theoretical derivation of kernel transformation for various data types (in 2.1), graphical network modelling (in 2.2) followed by results on simulated (in 3.1) as well as real life data (in 3.2).

## 2 | THEORY

In this paper, an unspecified source of information is highlighted as 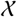, matrices are in capital bold (**X**), vectors in lower case bold (**x**), and scalars as italic lower case (*x*). Indices of matrices and vectors are highlighted as subscripts.

### 2.1 | Kernel representation

Kernels are useful methods for representing the population distribution as sample similarities. Let **K** be a symmetric *n* by *n* matrix indicating similarities between any pair of samples from an information source 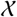 from a total of *n* samples. In the case of a matrix 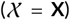, then the individual scalar elements of **K** can be defined as: **K**_*ij*_ = *f*(*x_i_, x_j_*) where f is a transformation function of the input vectors for sample *i* (*x_i_*) and *j* (*x_j_*). If a multivariate dataset of continous variables are assumed Gaussian, then the so called radial basis function kernelization (rbf-kernel) is a natural choice of transformation:

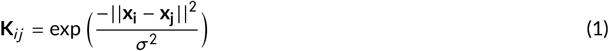

Where ||**x**_i_ – **x**_j_|| is the Euclidean distance and *σ*^2^ is the bandwidth of the kernel. For more information on kernel consult [2].

#### 2.1.1 | Data on survival-and recurrent episodes

Longitudinal information such as survival outcomes are characterised by a time-to-disease (*t* ≥ 0) and a disease status (zero if the disease has not occurred and one if it has occurred) within the time period of interest e.g. follow-up time from intervention or study period. However, in the case of data on recurrent episodes, a survival representation, merely reflects time to the first event, and thereby omits relevant information on remission and later occurring episodes. Be aware, that this representation is used for episodes of chronic conditions, such as asthma, where a diagnosis span from a few years to a lifetime. This is contrast to episodes of shorter duration such as common childhood symptoms, which is described in section 3.2 and further detailed in [8].

We propose a flexible kernel representation of longitudinal data defined as the sum of weighted kernels:

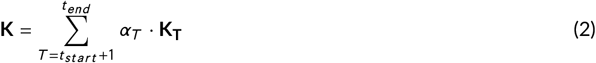

For classical survival data, where the disease status is binary, the similarities represented by the binary kernel: **K***_T=t_* at time *T* = *t* is defined as **K***_T=t_*(*i, j*) = 1 if *patient_i_* œ *patient_j_* (both have disease or both healthy) and *K*_T=t_(*i, j*) = 0 if not. This results in a sequence of binary kernels: **K_T=tstart_**, …, **K_T=t_end__**

The weights of each time kernel (*α_T_* ≥ 0) were defined as the change in values between two kernels at two consecutive time points, defined as the element wise change between two consecutive kernels:

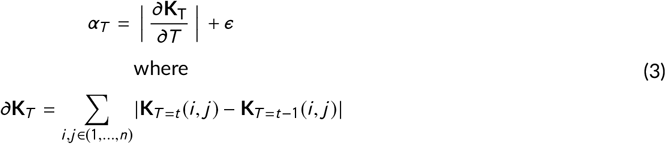

*ϵ* is a small number giving all periods a weight above zero and the first element of the weight vector is set to *ϵ* (*α*_1_ = *ϵ*). Further, the weights were scaled to a simplex (∑ *α_t_* = 1).

Traditionally survival analyses have been used for outcomes that are not reversible such as death. Over the years survival analyses have been applied to other types of data such as time to chronical diseases, time to relapse eg. Some patients go into remission after getting diagnosed with a disease. In order to capture this we introduce a third state to the model, the state of remission defined as healthy after a period of disease. Now we have three states: healthy, ill and remission and we define the similarity between two patients as:

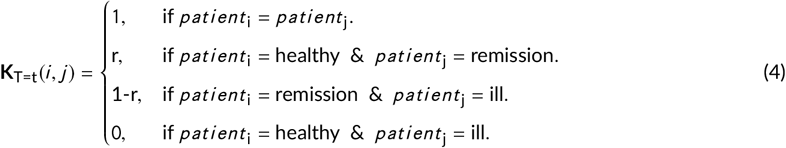

Where *r* is the rate of remission and **K_T=t_** ∈ {0, *r*, 1 – *r*, 1}. The weights and the final kernel were defined as in equation 3 and 2 respectively.

#### 2.1.2 | Repeated measurements of continuous data

The kernelization of data that does not follow data on time to event or recurrent episodes (as in section 2.1.1), but when applied to longitudinal continuous information such as repeated measurements of height or weight, the framework in equation 2 and 3 can be extended, by using a relevant measure of similarity for each time point (*T* = (*t*_1_, *t*_2_, …, *t_end_*)) such as:

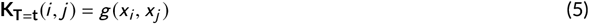

Where *g*(*x_i_, x_j_*) calculates a measure of similarity between *x_i_*, and *x_j_*: E.g. *g*(*x_i_, x_j_*) = exp(–(*x_i_* – *x_j_*)^2^/*s*) (with *s* being a relevant scaling constant).

The weights of the time kernels **K**_*T=t*_1__, **K**_*T*=*t*_2__, …, **K**_*T=t*_end__ are defined as in equation 3 and they are combined to Kas in equation 2.

### 2.2 | Graphical network based on partial correlations

#### 2.2.1 | Correlation coefficients

Kernel to Kernel correlations is defined via the so-called RV coefficient [5, 6], which simply amounts to a scalar correlation coefficient between two vectorized matrices. That is: The M individual kernels were unfolded, from **K**_*m*_ ~ (*n* × *n*) to vectors where the diagnoal elements are removed **k**_*m*_ ~ (*n*^2^ – *n* × 1). Each vectorized kernel was standardized to mean zero and standard deviation one. The correlation between two kernels can be expressed as in equation 6, which under centering and scaling (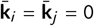 and *σ_k_i__* = *σ_k_j__*) reduces to an inner product. For detail on scaling of kernels and the definition of RV coefficient see Aben *et al* 2018 [1].

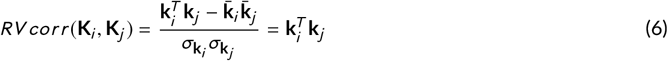

The correlations can then be combined in a correlation matrix **S** = **S**^*T*^ ~ (*M* × *M*).

#### 2.2.2 | Sparse Graphical LASSO

The matrix of unfolded standardized kernels is combined into a data matrix **X** = [**k**_1_, **k**_2_… **k**_*M*_] ~ (*n*^2^ – *n* × *M*), which is considered a manifestation of a multivariate distribution on what we term a *similarity manifold*. We use the GLASSO proposed by Friedmann et al 2009 [10] to estimate the partial correlations as the inverse covariance matrix on these standardized kernels, given the assumption that such a data representation can be subjected to a regularized maximum likelihood correlation estimation procedures as in [10].

The objective of the graphical lasso is.

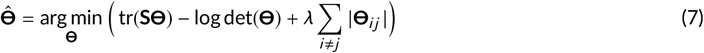

Where **S** = **S**^*T*^ ~ (*p, p*) is the observed covariance matrix, *λ* ≥ 0 is a tuning parameter and **Θ** = **Θ**^*T*^ ~ (*p, p*) is the inverse covariance matrix (or precision matrix).

In principal two algorithmic approaches have been suggested for solving this objective, either based on sequential updating of the covariance matrix or on the inverse of the covariance matrix (**Θ**) [11]. For the present work we utilize a dual optimization strategy for updating columns/rows of directly on **Θ** referred to as DP-GLASSO by [11]. Beyond superior convergence stability compared to GLASSO, this algorithm focuses directly on **Θ** which allows for an easy extension where certain parts of **Θ** are considered fixed (see section 2.2.3). For details on the derivation of the optimality criterion and the algorithm see Appendix A.

#### 2.2.3 | Introducing new variables to a fixed inner model

To be able to observe correlations between highly correlated biological data e.g. between different symptoms of illness (from here on the *inner network*) and not as highly correlated data such as environmental exposures or later disease without having to interpret to many ‘middle’ edges within the inner network, we propose a two-step approach. This method where the less influential biological correlations are removed offers simplicity in order to understand the complex biology. First, fit an inner network (**Θ_*inner*_**) of the biological data (here between different symptoms), followed by a model for the outer network conditioning on the inner correlation network, where the graphical structure both within the outer sources (**Θ_*outer*_**) of information as well as between the inner-and outer layers of data (**Θ_*inner2outer*_**) is estimated (see Figure 1 for a schematic representation).

**FIGURE 1.**
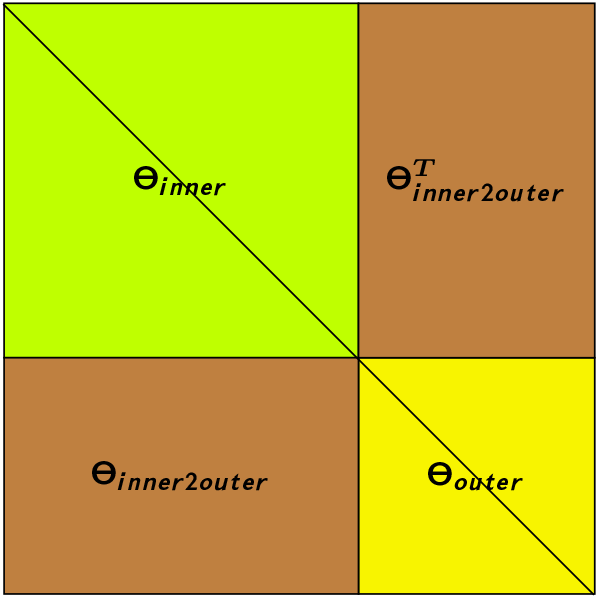
Adding an outer layer of data, and estimating a model (**Θ_*outer*_**, and **Θ_*inner2outer*_**) conditional on the *fixed* inner network (**Θ_*inner*_**).

This is achieved by a modification of the algorithm in A, such that the initialization of (**Θ**) is exchanged with the estimates from the inner network. Otherwise, the algorithmic procedure follows 1, with the only modification of *only* cycling through the columns associated with the new information.

### 2.3 | From graphical network to observation clustering/similarities

For the graphical network 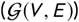 between with *M* nodes (|*V*| = *M*) the edge-set (*E*) is set by **Θ**. This network partitions information (here the M kernels) into clusters of correlations, however, without any explicit segmentation of the individual samples. Here, we propose a post-hoc procedure to derive a partition over the samples obeying the correlation structure in 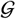.

First, for each of the *M* nodes, a network weight (*ω_m_*) is defined as the sum of the weights of its adjacent edges. That is:

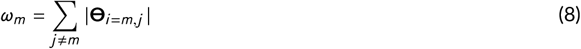

Hence *ω_m_* reflect how strongly node *m* is connected with the rest of the network. These weights are stored in a vector *ω* = (*ω*_1_, *ω*_2_, …, *ω_M_*). Using these weights, a *consensus kernel* (**KK**) can be derived as:

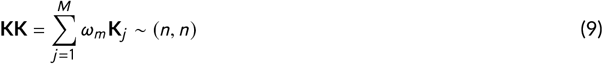

By definition, such a consensus kernel will reflect the population structure for the most connected nodes in the network, while disconnected nodes (*ω* = 0) will have no influence. By using hierarchical clustering on this consensus kernel a partition of the n samples, reflecting the dominating correlation patterns in the data, will be derived.

#### 2.3.1 | Model selection

##### Penalty sequence

Depending on the problem, the *dynamic* sequence of penalties *λ_min_*, …, *λ_max_* is not trivial to set, which essentially results in screening sequences of models for penalty values with no practical change in parameter estimates. For this reason the *dynamic* penalty sequence is set such that *λ_min_* results in a single off-diagonal element of **Θ** being 0, and similarly *λ_max_* returns *only* a single active (≠ 0) off-diagonal element.

##### Model section criterion

Cross-validation evaluating the non-regularized maximum-likelihood based on a trained precision matrix 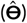 in comparison with an independent (left-out) observed covariance matrix 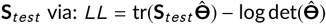 is generally good for *overall* control of estimation flexibility. However, this procedure suffers from being unable to correctly identifying 0 elements in the precision-matrix. This is due to shrinking these parameters to a small non-zero neglect able entity, that have no practical influence on the cross validation fit, but leads to a false discovery. This phenomena is previously described by Weijie *et al* 2017 [12]. For this reason a criterion explicitly weighting the amount of non-zero elements has been proposed via the extended Bayes Information Criterion [7], which penalizes dense graphs with increasing *γ*, following:

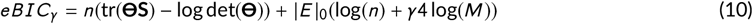

where *n* and *M* is the number of samples and number of graph nodes (kernels) respectively, |*E*|_0_ is the number of edges. I.e. the number of non-zero of diagonal elements of **Θ**, and *γ* is a tuning parameter putting emphasize on finding the zero elements of **Θ**.

## 3 | RESULTS

### 3.1 | Simulations

For this part we pertain to the individual data being low rank multivariate data, simulated as bilinear matrices. The block-wise dependency is constructed by a linear mapping of sample related component scores in one block to another block. On top of the systematic bilinear structure there is added either structured noise or white noise at varying levels.

For this setup we construct five blocks (**X**_1_, **X**_2_,… **X**_5_) of each *n* samples and *p*_1_, *p*_2_,… *p*_5_ variables following the structure shown in figure 2.

**FIGURE 2.**
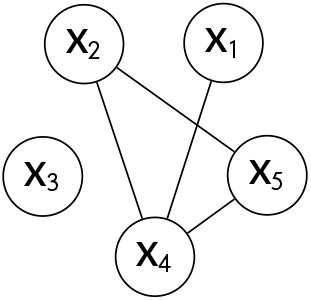
Structure of a five block network with a disconnected or *isolated* vertex (*degree* = 0, **X**_3_, also known as an independent set), a *leaf vertex* (*degree* = 1, **X**_1_) and cyclic connected vertexes (**X**_2_, **X**_4_, **X**_5_)

We define the scores for the five blocks as in equation 11 and 12:

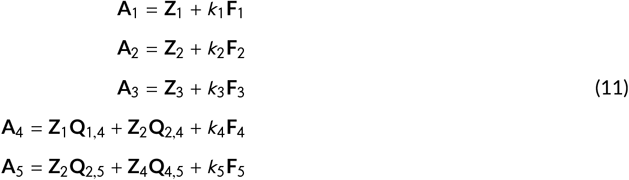

The matrices **Q**_*i,j*_ constitutes the mapping between block *i* and *j*, and **F**_*i*_, constitutes the *structured* noise, where the levels is set by ***k**_i_*. The matrices **Z**_1_ ~ (*n, c*_1_), **Z**_2_ ~ (*n, c*_2_), **Z**_3_ ~ (*n, c*_3_), **Q**_1,4_ ~ (*c*_1_, *c*_4_), **Q**_2,4_ ~ (*c*_2_, *c*_4_), **Q**_25_ ~ (*c*_2_, *c*_5_), **Q**_45_ ~ (*c*_4_, *c*_5_) and **F**_1_ ~ (*n, c*_1_),…, **F**_5_ ~ (*n, c*_5_) all have entries drawn from a Gaussian distribution 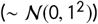. The rank (*c_i_*) of the individual data blocks are drawn from a uniform distribution 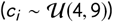.

The five data sets are now given by:

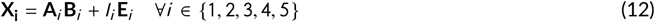

where **B**_*i*_, ~ (*c_i_*, *p_i_*) is the right component loading matrix drawn from a Gaussian distribution 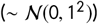. The number of variables (*p_i_*) for each data block is drawn from a uniform distribution 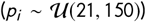. The noise level scalars *k_i_*, and *l_i_*, are set to fulfill signal to noise levels for i) structured noise defined as:

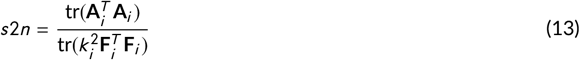

as well as for ii) unstructured white noise defined as:

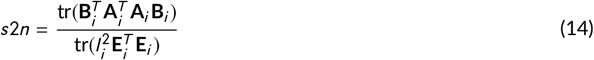

The data sets **X**_1_, **X**_2_, …, **X**_5_ are kernelized such that *K_i_*, ~ (*n, n*) is the kernelization of **X**_*i*_ ~ (*n, p_i_*) using the exponential of the negative euclidean distance between pairs of observations.

We investigate the effect of number of samples (*n* ∈ (20,50,100,200,500)), type of noise (*White* or *Structured*), signal to noise (*s*2*n* ∈ (1/8,1/4,1/2,1, 2,4, 8)) on the ability to recover the underlying structure represented as sensitivity (true positive rate), specificity (true negative rate) and the combination of the two as the area under the receiver operator characteristics curve (AUC). For all simulationsmodel selection is done using eBIC.

For each of the 700 design combinations 100 simulations are produced, and model selection is done by eBIC with *γ* ∈ (0,0.5,1).

Figure 3 shows the performance in terms of true positive- and true negative rate as well as the agglomerated area under the receiver operator curve (AUC). These quality metrics are shown for models selected using the eBIC with *γ* ∈ (0,0.5,1). Further, included is also the *oracle* model, which is the model that most optimally resembles the underlying edge set (figure 2) in terms of AUC. The behaviour of the true positive rate follows general statistical estimation paradigm with higher *n* and signal to noise yielding more accurate results. However, this is at the cost of the true negative rate which declines with these data characteristics. In general, the difference between the *oracle* model and the model selected using eBIC is that *λ_oracle_* - *λ_eBIC_* is biased positively for low signal-to-noise and low sample size, turning into being biased negatively with high signal-to-noise, reflecting that eBIC is too conservative when dealing with low-noise data, and too optimistic in the case of small-and noisy data. (see Appendix B for details)

**FIGURE 3.**
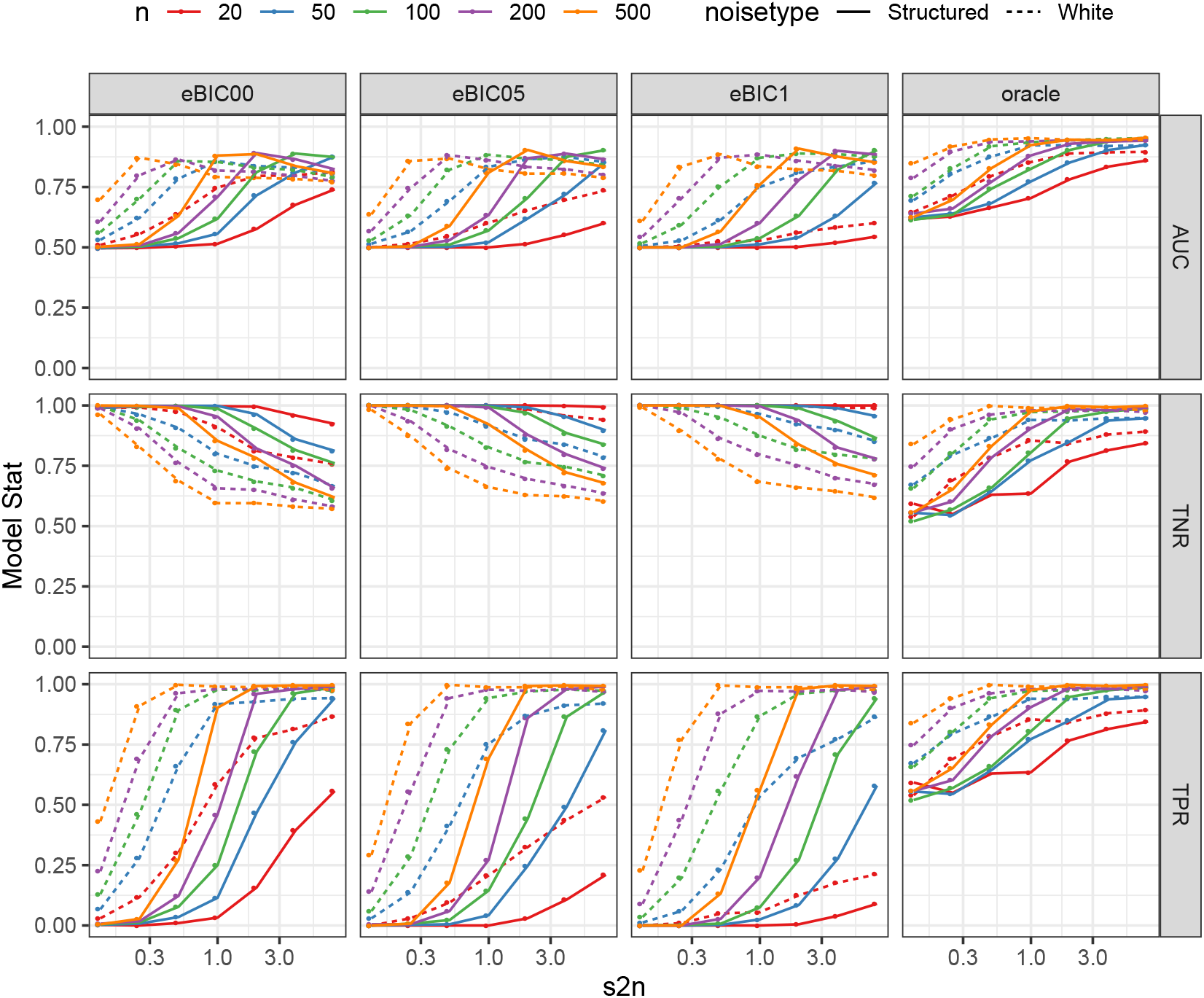
Area under the receiver operator curve (AUC), True Negative Rate (TNR) and True Positive Rate (TPR) for simulated data with varying signal to noise (*s2n* - x-axis), number of samples (*n* colors) and type of noise (line type). The best model is selected based on eBIC with *γ* ∈ (0,0.5,1) as well as - the unobserved - *oracle* (defined as the model with the best AUC). Each point is the mean of 100 simulations.

Figure 4 investigates from which parts of the graph the uncertainty derives. Here, node **X**_3_ is the single isolated / non-connected vertex of the graph, and it seems relatively easy for the model to also correctly exclude edges to this node. The indirect connection between **X**_1_ and **X**_2_ and **X**_5_ (via **X**_4_) is causing the observed sub optimal true negative rate. For the identification of the true connections, generally similar results is observed. There seem to be no discrepancy in the estimation power depending on whether the edge is in a fully connected part of the network (between **X**_2_, **X**_4_ and **X**_5_), or towards the leaf vertex **X**_1_ from **X**_4_.

**FIGURE 4.**
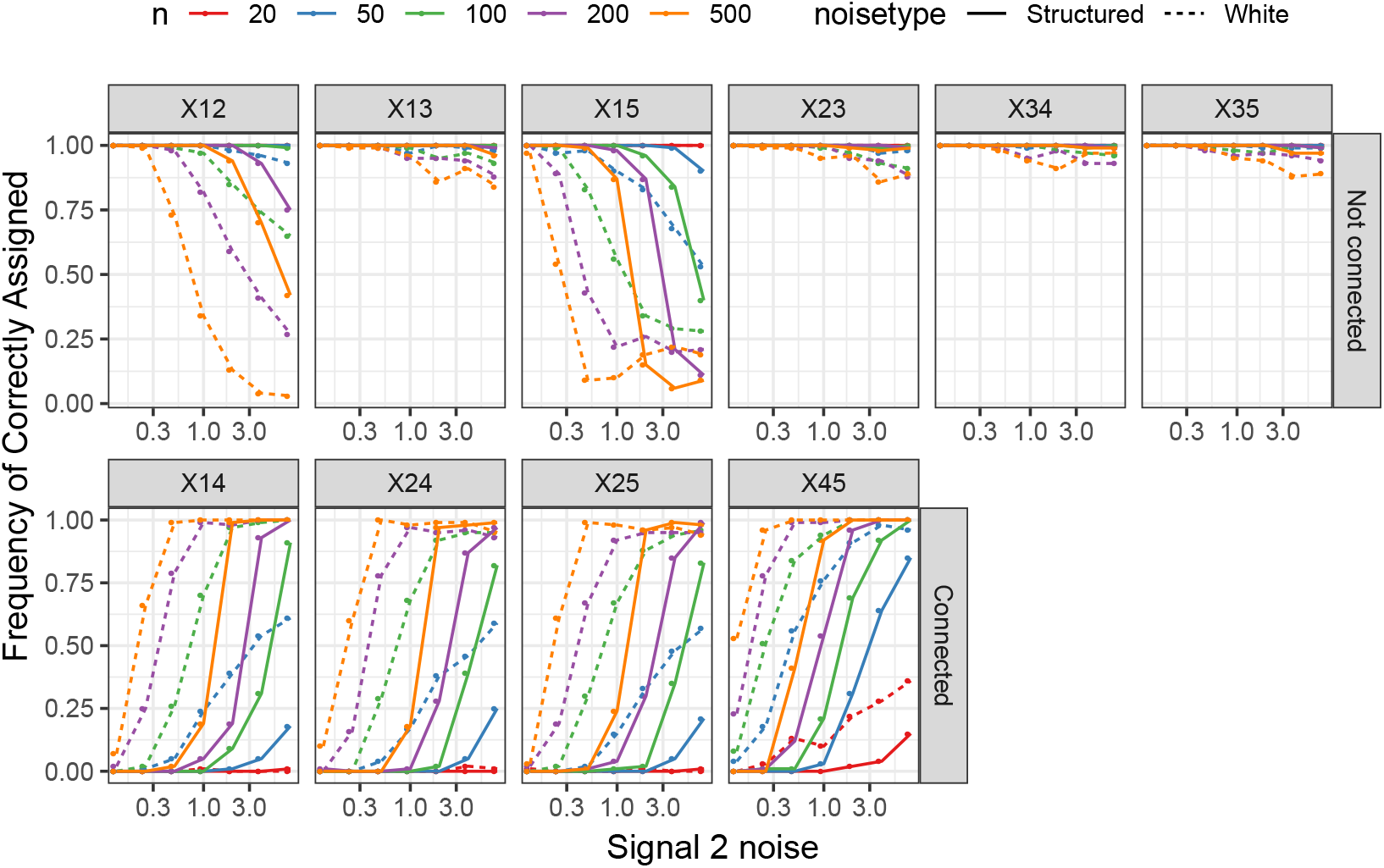
For each of the 10 possible connections (4 positive and 6 negative) the frequency of correct identification is computed, using the model selected by eBIC with *γ* = 0

### 3.2 | COPSAC symptom burden

From a total of *n* = 403 children, with at least 50% of diary registration each month, the symptom burden was prospectively registered daily the first three years of life, where the following common childhood symptoms were registered: *cough, breathlessness, wheeze, cold, pneumonia, inflammation of the throat, ear infection, fever, gastric infection* and *eczema*. For details on these data including seasonal variation see [8]. For this analysis, the symptoms of the 2^nd^ and 3^rd^ year were investigated, and further were the symptoms *breathlessness* and *wheeze* combined in to one symptom due to their similar nature. This gives nine symptoms over two years resulting in 18 symptom kernels.

Additionally, 21 exposures obtained during or prior to the first year as well as the asthma-and eczema status up to age 6 years were incorporated. Of particular interest is to answer **Q**_1_: How early life exposures leads to later development of asthma? and here; **Q_2_**: How the early life symptom burden can be considered as a path towards later disease.

#### 3.2.1 | Kernelization of symptom diaries

To reduce noise, the daily diary registrations were summarised (fractions of days with the symptom out of registered days) for each calendar month the 2^nd^ and 3^rd^ year of life (January to December, year 2 to year 3), resulting in 24 registrations per symptoms per child. The distance between any two children was constructed using the Euclidean distance between frequency vectors within each year. By explicitly factoring in calendar month in the construction of the pairwise differences, results in kernels that allow one to disentangle seasonal symptoms patterns. For instance, two children with an overall equal prevalence but with distinct seasonal profiles, will be rather dissimilar in the kernel. The distance measures were transformed to measures of similarities by multiplying by minus one. For details on the choices related to kernel transformation see Appendix C a well as [8].

#### 3.2.2 | Graphical inner-model

The 18 kernels of symptoms were unfolded, the diagonals were removed and they were standardized with mean zero and standard deviation one. Model selection were performed using the eBIC with *γ* = 0 based on a 10 times repeated 5-fold cross-validation. Figure 5 shows the cross-validation results, where a penalty of *λ* = 0.076 was chosen resulting in 118 edges (out of 153 possible - 77%).

**FIGURE 5.**
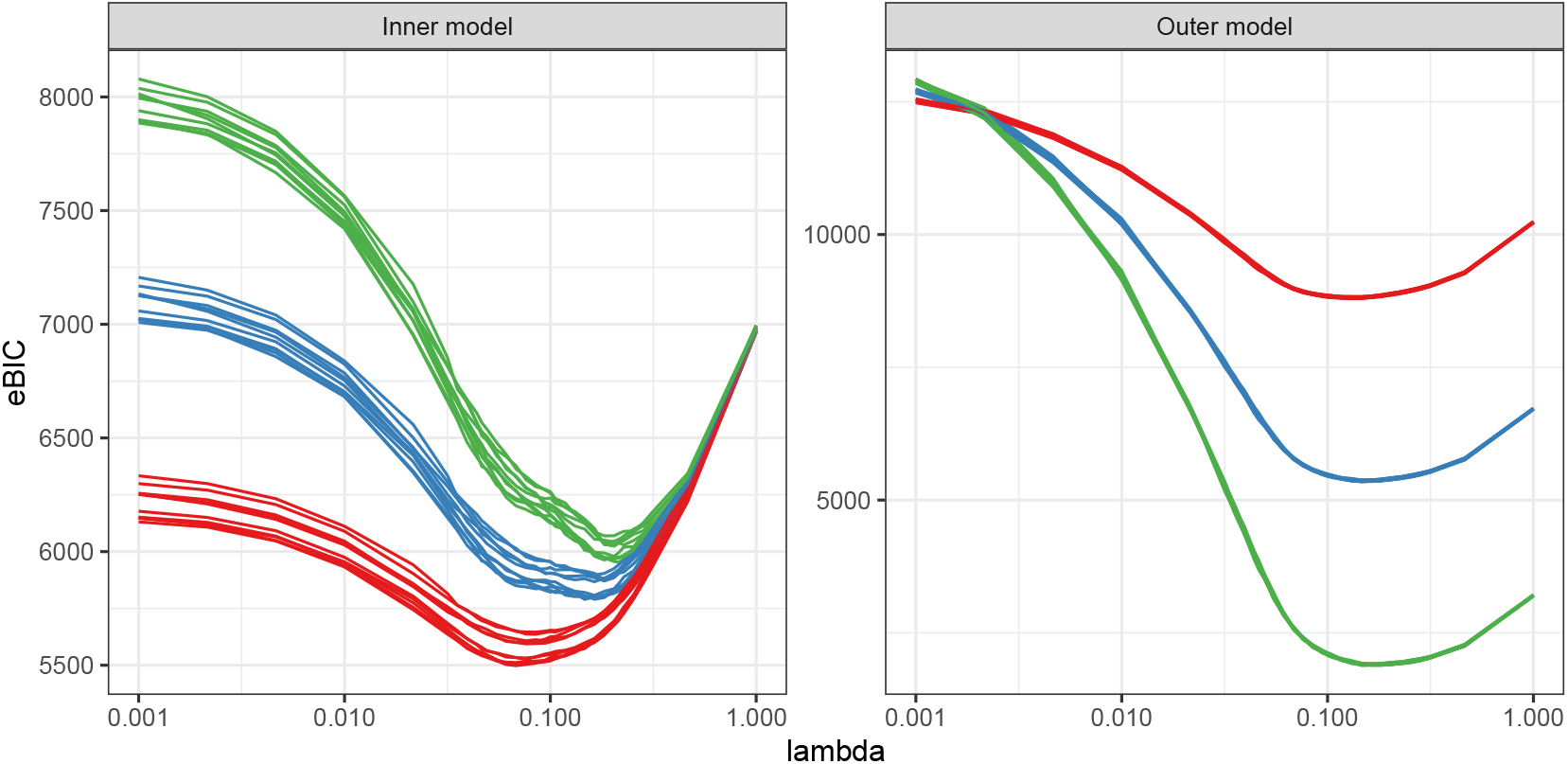
10 repetitions of 5-fold cross validated GLASSO models with eBIC as model selection criterion. Color refers to y value of eBIC (red: *γ* = 0, blue: *γ* = 0.5, green: *γ* = 1). Panels relates to the *inner* and *outer* models

Table 1 shows the number of connections for each symptom-year combination. The node with the most connections (ten) is fever 2^nd^ year, while eczema 3^rd^ year was only connected to eczema-and cough 2^nd^ year.

**TABLE 1.** Node characteristics for the network model in terms of number of edges in total (*Edges*) as well as towards symptoms (*E2symptom*), exposures (*E2exposure*) and disease (*E2disease*). Only connected nodes are listed

|  | Year | Symptom | Edges | E2symptom | E2exposure | E2disease |
| --- | --- | --- | --- | --- | --- | --- |
|  | 2 | BW | 10 | 8 | 1 | 1 |
|  | 2 | Cold | 7 | 6 | 0 | 1 |
|  | 2 | Cough | 9 | 7 | 1 | 1 |
|  | 2 | Ear | 8 | 8 | 0 | 0 |
|  | 2 | Eczema | 8 | 7 | 0 | 1 |
|  | 2 | Fever | 10 | 10 | 0 | 0 |
|  | 2 | Gastric | 6 | 6 | 0 | 0 |
|  | 2 | Pneumonia | 6 | 6 | 0 | 0 |
|  | 2 | Throat | 5 | 5 | 0 | 0 |
|  | 3 | BW | 8 | 6 | 1 | 1 |
|  | 3 | Cold | 10 | 9 | 0 | 1 |
|  | 3 | Cough | 12 | 9 | 2 | 1 |
|  | 3 | Ear | 4 | 4 | 0 | 0 |
|  | 3 | Eczema | 3 | 2 | 0 | 1 |
|  | 3 | Fever | 8 | 8 | 0 | 0 |
|  | 3 | Gastric | 4 | 4 | 0 | 0 |
|  | 3 | Pneumonia | 9 | 9 | 0 | 0 |
|  | 3 | Throat | 4 | 4 | 0 | 0 |
| exposure |  | Cat birth | 1 | 0 | 1 | 0 |
| exposure |  | Daycare type | 1 | 0 | 1 | 0 |
| exposure |  | Daycare age | 1 | 0 | 1 | 0 |
| exposure |  | Dog birth | 1 | 0 | 1 | 0 |
| exposure |  | Gestational age | 3 | 1 | 2 | 0 |
| exposure |  | Maternal asthma | 3 | 2 | 1 | 0 |
| exposure |  | Maternal BMI | 2 | 0 | 2 | 0 |
| exposure |  | Maternal rhinitis | 1 | 0 | 1 | 0 |
| exposure |  | Paternal asthma | 1 | 0 | 1 | 0 |
| exposure |  | Paternal rhinitis | 1 | 0 | 1 | 0 |
| exposure |  | Preeclampsia | 1 | 0 | 1 | 0 |
| exposure |  | Premature | 4 | 1 | 3 | 0 |
| exposure |  | C-section | 2 | 0 | 2 | 0 |
| exposure |  | Smoking 3<sup>rd</sup> trimester | 1 | 1 | 0 | 0 |
| disease |  | Asthma 6yrs | 6 | 6 | 0 | 0 |
| disease |  | Eczema 6yrs | 2 | 2 | 0 | 0 |

#### 3.2.3 | Graphical outer-model

Data on exposure (*Cat at birth, Dog at birth, Daycare type, Daycare age, Gestational age, Maternal asthma, Maternal BMI, Maternal rhinitis, Paternal asthma, Paternal rhinitis, Preeclampsia, Premature delivery* and *Mode of delivery*) was kernelized and treated in the same way as the symptom kernels.

The asthma diagnoses were kernelized as proposed in section 2.1.1. 85 (21 %) of the children got a diagnosis of asthma before the age of six years and 61 (72%) went into remission before the age of six years.

123 (31%) of the children got a diagnosis of eczema before the age of 6 years. Of these, 77% (95 children) went into a state of remission at the age of 6 years, and had thereby lost this diagnosis. Due to the early debut of eczema peaking around age 1-2 years, the eczema kernel were constructed based on those having the diagnosis *cross sectional* at age 6 years. This in order to avoid correlations by construction as symptoms of eczema is also included in the diary registrations.

These information sources, in the form of kernels, were added to the symptom burden data, to establish a model between risk factors, early life symptoms and chronic diseases. The partial correlations between symptoms found in section 3.2.2 were fixed in this model.

The outer partial correlation network was found with the customized conditional GLASSO, as proposed in section 2.2.3. The outer model were selected at *λ* = 0.1 based on 10 times repeated 5-fold cross validation using eBIC with *γ* = 0 as criterion. The corresponding network consisted of additional 22 edges of which 5 were between symptoms and exposures, 8 between symptoms and disease and 9 within exposures (see Table 1 for summary details of the graph). The resulting model is shown in figure 6.

**FIGURE 6.**
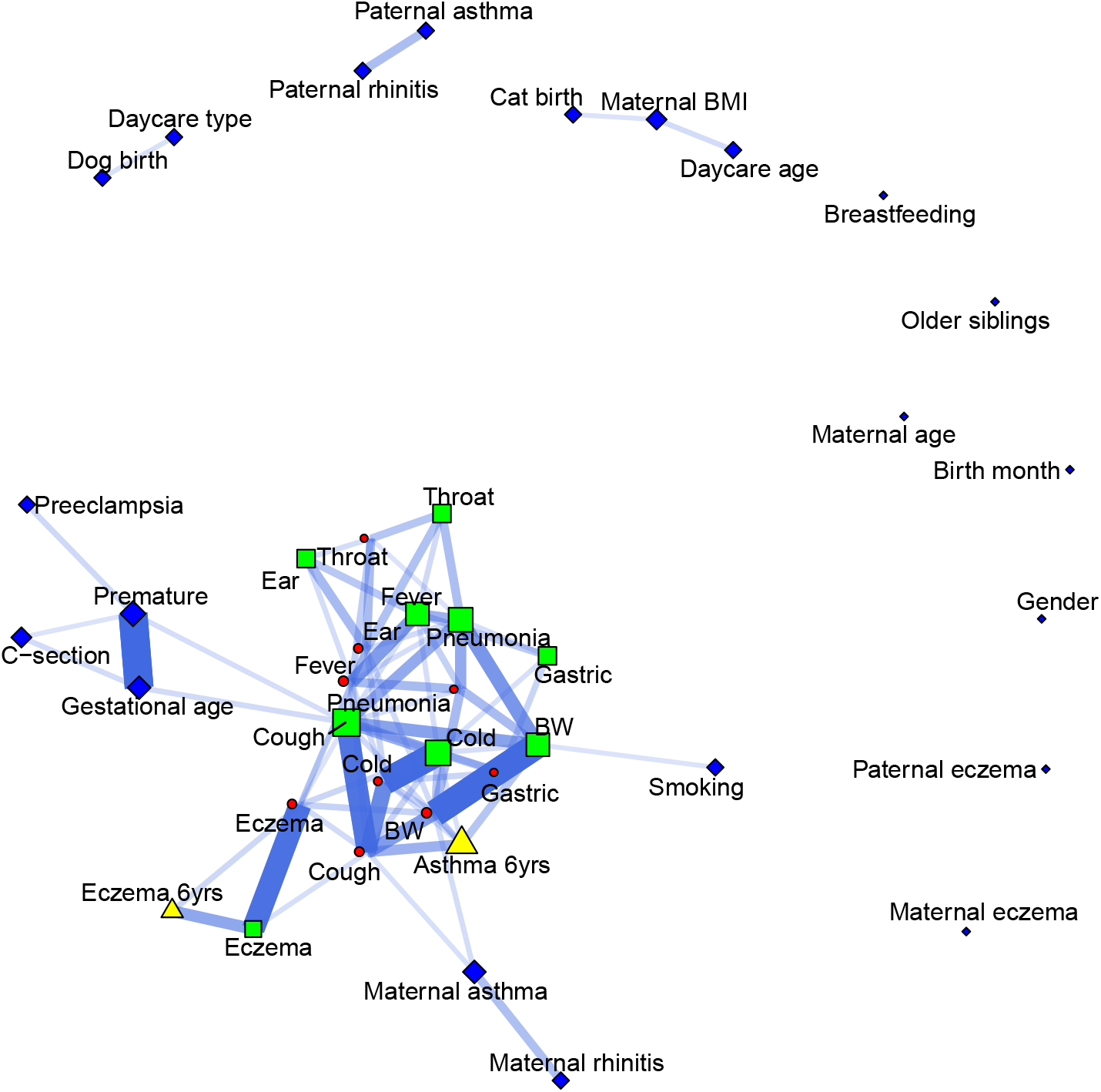
Partial network of symptoms in the 2^nd^- (red circles) and 3^rd^ (green squares) year of life in relation to exposures (blue diamonds) and diseases at age of six years (yellow triangles). The edge thickness and opacity indicates strength of relation, and the node size indicates the network centrality for the individual node

The airway symptoms: cough, breathlessness-wheeze and cold are clustered together in the center of the graph with connections to a cluster of eczema (year two, three and cross sectional at age six years) and a cluster of the rest of the symptoms (ear and throat infections, fever, pneumonia and gastrointestinal infections) on the other side. These symptoms are generally thought to be driven by bacterial or viral infections [13]. Maternal asthma is connected to cough and breathlessness-wheeze the second year of life. Pregnancy-and birth circumstances represented by gestational age and prematurity is connected to cough the third year, and interestingly the well known risk factor Cesarean section for a wide range of diseases [14], is only related to early life symptom burden through early delivery. Furthermore, maternal smoking during third trimester of pregnancy is connected to breathlessness-wheeze third year of life in line with previous findings [15]. These connections suggest an environmental effect on the respiratory health in early life, where as for the symptoms grossly driven by infections, the connections with risk factors are indirect.

#### 3.2.4 | Clustering children based on the inner-model

The proposed graphical model estimates clusters of associated information. However, it can be fortunate to also group the children into certain groups based on their symptom burden pattern as a phenotypic characterization. Based on the inner network between nine symptoms across year 1 and 2 a consensus kernel is calculated using the sum edge-weights for each node (see table 1). Hierarchical clustering results in a dendrogram as shown in figure 7, which is cutted to partition the children into 3 clusters with a distribution of ((*n_red_* = 12, *n_green_* = 26, *n_blue_* = 365)), where the small *red* cluster corresponds to the most diseased children followed by the *green* cluster, leaving the large *blue* cluster being the overall most healthy.

**FIGURE 7.**
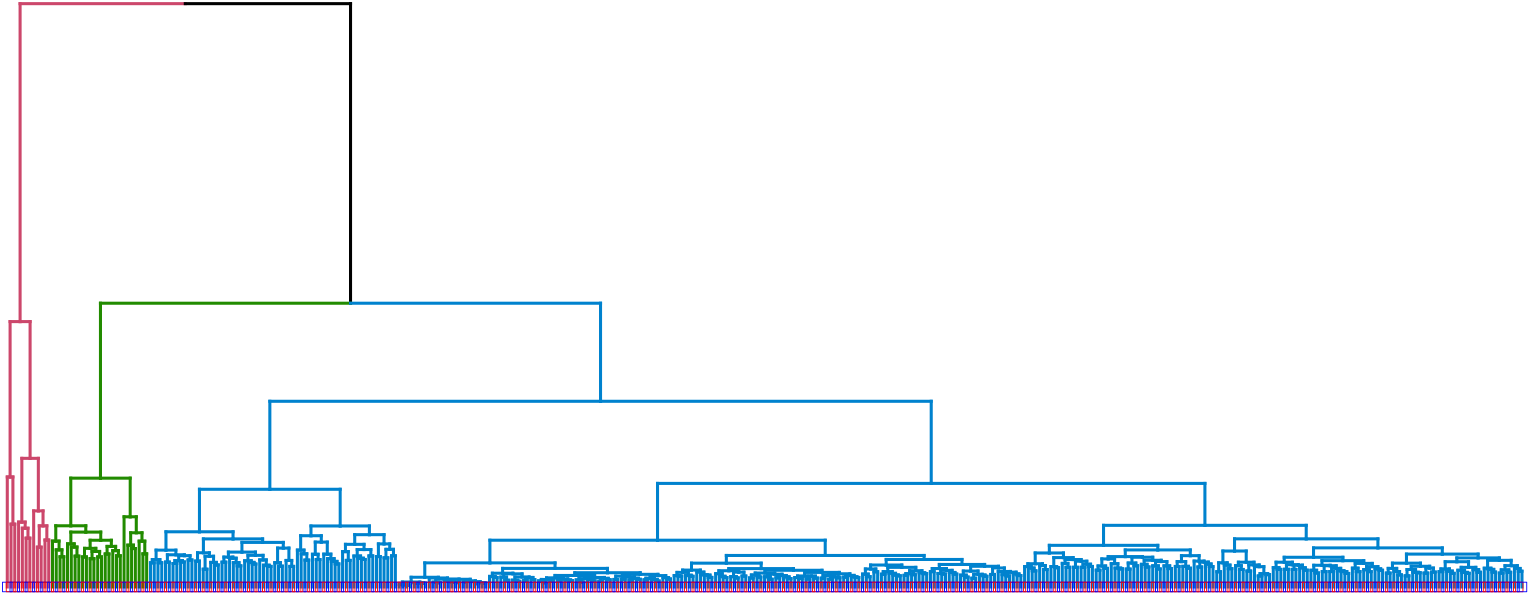
Dendrogram of consensus kernel based on symptom network splitted into 3 clusters with a distribution of (*n_red_* = 12, *n_green_* = 26, *n_blue_* = 365)

## 4 | DISCUSSION

We propose a flexible framework to do explorative analyses of heterogeneous data encompassing longitudinal symptom observation, survival type disease data and various data on exposure. The method is based on kernel transformation for data harmonization followed by graphical network modelling by the graphical LASSO to uncover direct and in direct correlations.

Kernels are flexible and can capture differences shifted in time, on different scales and of distributional nature. However, how the original data is kernelized will have a large impact on which patterns that will be captured. In particular for this work we have chosen to factor in calendar month in the transformation of data, due to the fact that there is a large seasonal component in symptom burden where for instance eczema is more prevalent during the changing seasons (autumn and spring) and respiratory symptoms are more prevalent during the winter months (for details on symptom burden seasonal patterns see [8]). In detail, consider two children (A and B) with the same overall symptom burden prevalence, but where child A have episodes during winter, where as child B have the episodes evenly spread across the year. By using the proposed kernel transformation, such differences will be conserved, and possibly affect the GLASSO model.

A dual representation of the well-known GLASSO were used for graph estimation. This dual formulation works directly on the entity of interest; the precision matrix, as opposed to the standard GLASSO algorithmicly working on the covariance matrix. Beyond numerical stability, this allows to fix certain parts of the precision matrix via a very simple modification of the algorithm.

The framework is validated on both simulated and biological data. The simulations show that capturing true edges follows a normal statistical paradigm with better recovery for larger *n* and *signal-to-noise*, and further that unstructured *white noise* interfere less with the model estimation than structured noise. However, capturing non-existing edges behaves oppositely, with a higher false negative rate with larger *n* and *s*2. This phenomenon derives from indirect relations, and not from spurious edges towards *disconnected* nodes (edges towards **X**_3_ in Figure 4). Overall though, the error-rate in terms of AUC increases with *n* and *s*2*n*.

The integration of the symptom burden data during 2^nd^ and 3^rd^ year of childhood showed that the respiratory symptoms are highly correlated, and that the ear infection and the inflammation of the throat are connected which corresponds with the bacteria moving from the throat to the ears in line with the coherent airway epithelium in these communicating organs [13].

Introducing precursors of disease as well as asthma and eczema by age 6 years to the symptom network highlights that maternal asthma is connected to respiratory symptoms in the 2^nd^ year of life, while maternal rhinitis only associates to respiratory symptoms indirectly through maternal asthma. Further, maternal asthma is suggested to be a risk factor for the development of asthma by age 6 years indirectly through an elevated level of respiratory symptoms in early life. Paternal asthma, rhinitis and eczema do not have any connections to any of the symptoms. This corresponds with previous research showing maternal asthma increases the risk of childhood asthma to a higher degree than paternal asthma [16,17].

Networks behave different based on their correlation structures. When only investigating the correlations between the symptoms, we observe a network with many well-connected nodes. When exposures/environmental factors are included in this part of the network has a weaker correlations between symptoms and exposures with fairly many (7) disconnected nodes. Furthermore, we see that few exposures (4) associate directly with the symptoms. However, the pertinent challenge here, is to distinguish between associations of symptoms which are timely matched and are having overlapping phenotype presentation with those of exposures or disease, which are disparate in time and a-priory of much less strength. By using the two step estimation procedure with an *inner* model on symptom data only, followed by an *outer* model on all data conditioning on the inner estimates allows for using a higher penalty on the inner model (*λ_inner_*) to only include *strong* connections, while allowing the penalty of the outer model (*λ_outer_*) to include everything that generalizes in terms of cross-validation.

## 5 | CONCLUSIONS

A combination of kernel transformation and graphical LASSO is proposed for integration of heterogeneous data layers. The method is demonstrated on both simulated-and real life data pertaining to early life symptom burden, precursors and development of disease at age 6 years.

## 6 | SOFTWARE

All algorithms have been coded in Matlab ®R2019 and is available through github: (https://github.com/mortenarendt/KerGLASSO), the code depends on the quadprog() function from the optimization toolbox(http://www.mathworks.com/products/optimization/). Plotting is conducted using R(ver 3.5.1) with `ggplot2, tidygraph` [18], `ggraph` [19] and `igraph` [20] for graph handling and plotting.

## Acknowledgements

We thank the children and families of the COPSAC cohort study for all their support and commitment. We acknowledge and appreciate the unique efforts of the COPSAC research team.

## A | DERIVATION OF DP-GLASSO

Let 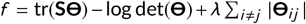 be the objective from equation 7. And further, let **W** = **Θ**^−1^ be the population covariance- and precision matrix respectively, and **S** be the observed covariance matrix. All symmetric and of size (*p, p*).

The sub-gradient of *f* with respect to **Θ** can be formalized as:

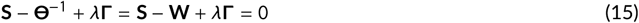

Where **Γ** is the sign matrix of the individual elements of **Θ**. Let:

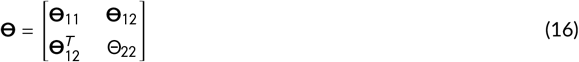

be a representation of **Θ** where **Θ**_11_ ~ (*p* - 1, *p* – 1), **Θ**_12_ ~ (*p* – 1,1) and Θ_22_ ~ (1,1) (with a similar representation for **S** and **W**). Using calculus for matrix inversions reveals the block wise stationary point of equation 15 to be:

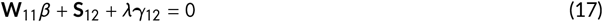

where *β* = **Θ**_12_/Θ_22_, and *γ*_12_ is a sign vector of **Θ**_12_. This linear system of equations 17 is the score (or normal-) equations associated with the LASSO with the following quadratic objective:

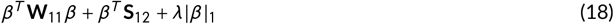

The dual representation of this problem (equation 18) is derived by defining a new vector: **x** = *β* and extending the objective to:

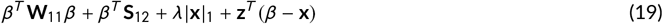

where *z* is a vector of (non negative) Lagrangian multipliers.

The stationary points of objective 19 with respect to *β*, assuming **W**_11_ ≻ 0, is:

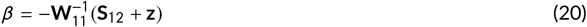

and with respect to **x**

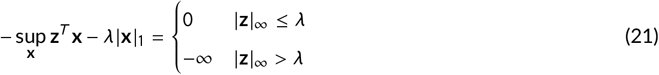

Combining (19), (20) and (21) leads to a dual quadratic minimization problem with a box constraint on the Lagrangian multipliers:

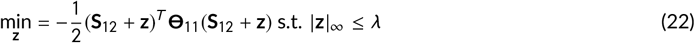

The primal estimates follows from equation 23

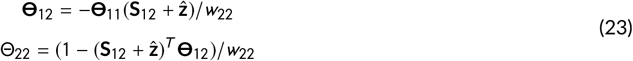

Without loss of generality, the entire solution can be achieved by rearranging the columns and rows of **S, W** and **Θ** to cycle through all columns of the covariance-and precision matrix with sequential conditional updates. See algorithm 1 for the entire procedure.

**Algorithm 1:**
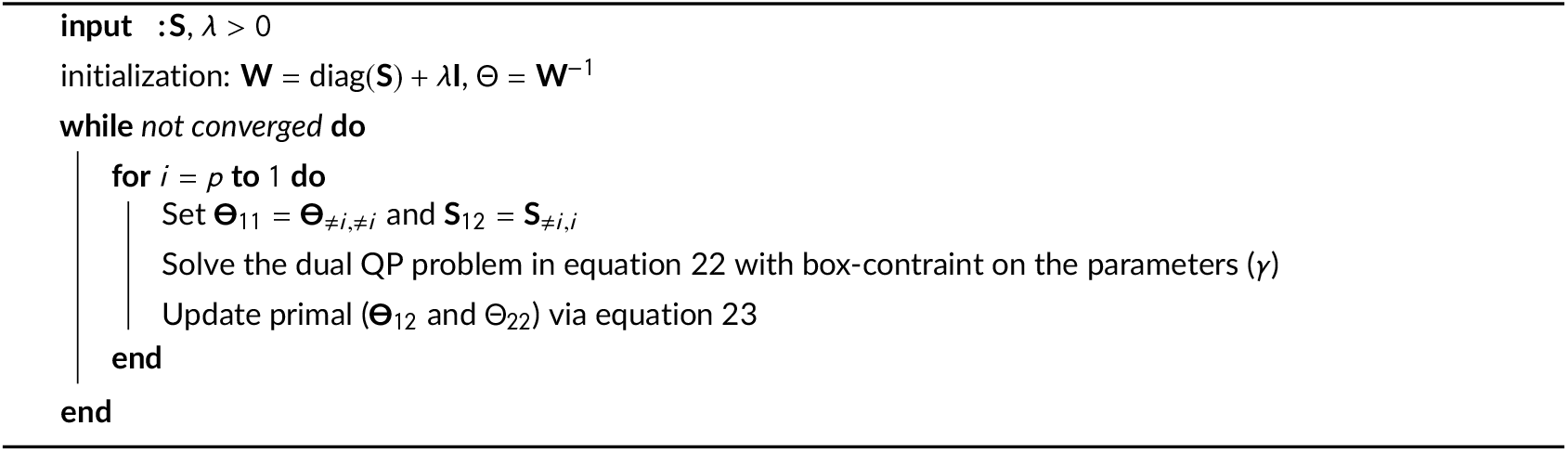
DP-GLASSO

## B | MODEL SELECTION - SIMULATION STUDY

The behaviour in the classification accuracy in Figure 3 and 4, especially when contrasting model selection with the oracle is due to a biased hyper parameter selection (λ) depending on the sample size and level of noise.

Figure 8 show this behavior when comparing the hyper parameter for the oracle model with that selected by eBIC.

**FIGURE 8.**
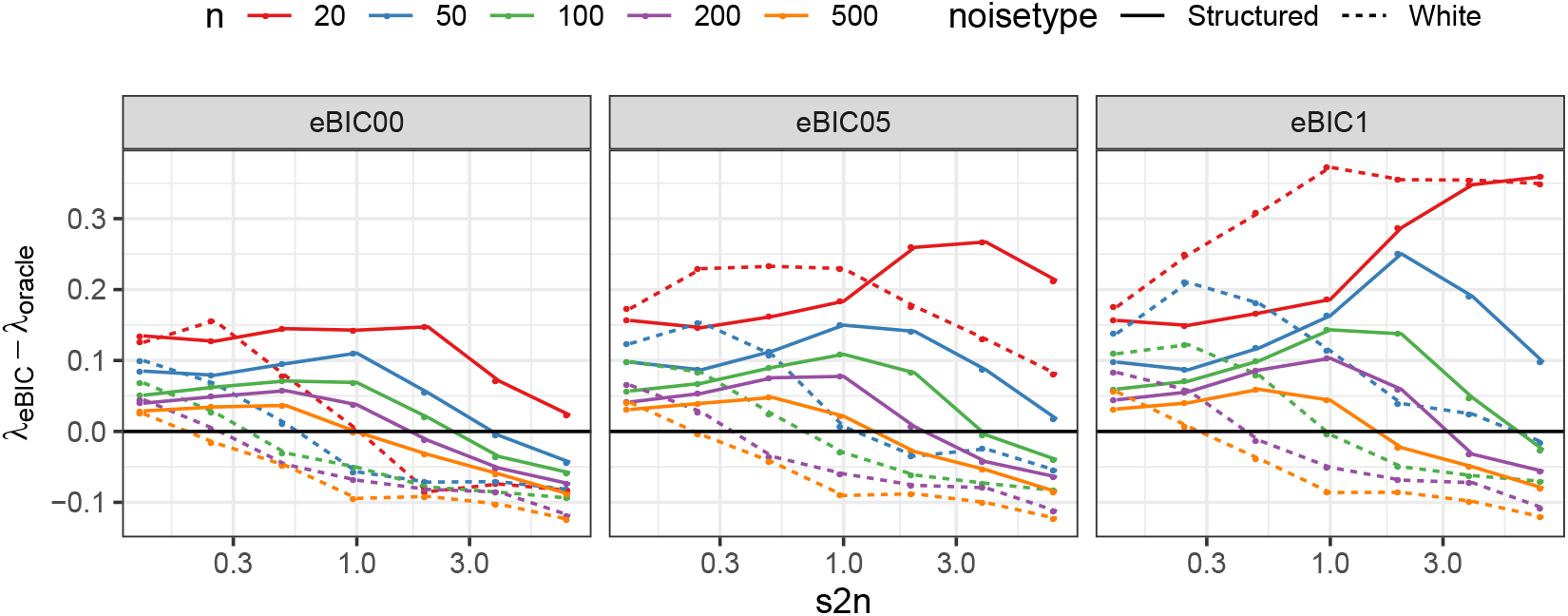
Hyper parameter difference between oracle model and model based on eBIC, as function of signal to noise (*s2n* - x-axis), number of samples (*n* colors) and type of noise (line type). Each point is the mean of 100 simulations.

## C | KERNEL TRANSFORMATION OF SYMPTOM BURDEN DATA

The raw diary symptom burden data represents binary (whether or not) the child had symptoms. This, for each day across a total of nine symptoms.

In order to investigate the effect of 1) time resolution, 2) distance metric and 3) distance to similarity function, a design of experiment were established.

The factors were:

### Biological

- **Age of child** (#2) second year or third year
- **Symptom** (#9) breathlessness or wheeze, cold, cough, ear infection, eczema, fever, gastric, inflammation and pneumonia

### Technical

- **Time-resolution** (#4) The symptom burden is agglomerated over calendar-– *day* (1,… 365)
  – *week* (1, …, 52)
  – *month* (1, …, 12)
  – *year* (total prevalence) into the number of days with symptoms within each bin.
- **Distance metric** (#4) Euclidean, City block or Manhattan, Hamming and Hamming on presence/absence data
- **Distance to similarity** transformation (#2) as 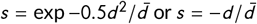, where *s* and *d* are measures of similarity and distance respectively, and 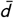 is the average distance over the entire (*n, n*) kernel of distances: 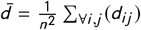.

This result in a 32 different options for each symptom and year combination, leading to a total of 576 kernel realizations. These were compressed by tucker3. Figure 9 shows the first two component scores. Within the upper 8 panels (and the lower 8 panels) only technical choices influence the distribution. In principal, The largest discrepancy between scores were due biological differences (symptoms and child age) while the choice of distance metric (reflected by color) and the distance to similarity function only contribute to minor differences. Agglomeration over day, week and month reveals similar patterns, while truncation of the information into a single yearly prevalence makes the results less coherent in terms clustering of symptoms.

**FIGURE 9.**
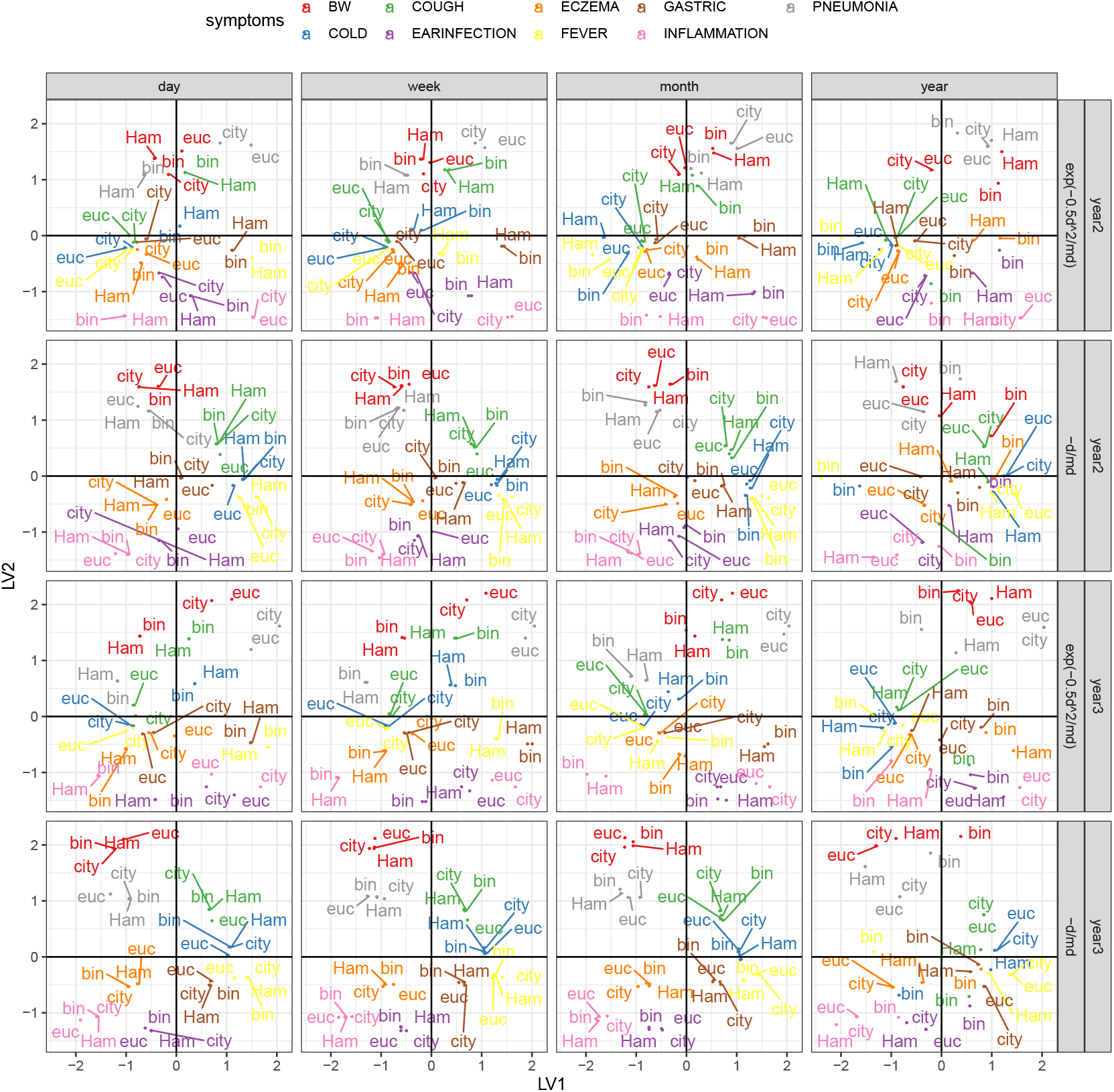
Component 1 and 2 from a tucker3 decomposition of 576 different representations of the 9 by 2 (symptoms by age). The panels are row wise: age of child and distance to similarity transformation and column wise: time resolution into comparing calendar-day, week, month and year. Colors reflects symptoms, while label reflects distance metric (*euc = Euclidian, city = city block, Ham = Hamming* and *bin = Hamming on binarized data*)

## references

[1] Aben N, Westerhuis JA, Song Y, Kiers HA, Michaut M, Smilde AK, et al. iTOP: inferring the topology of omics data. Bioinformatics 2018;34(17):i988 –i996.

[2] Shawe-Taylor J, Cristianini N, et al. Kernel methods for pattern analysis. Cambridge university press; 2004.

[3] Camps-Valls G, Gómez-Chova L, Muñoz-Marí J, Rojo-Álvarez JL, Martínez-Ramón M. Kernel-based framework for multitemporal and multisource remote sensing data classification and change detection. IEEE Transactions on Geoscience and Remote Sensing 2008;46(6):1822–1835.

[4] Carroll JD, Chang JJ. Analysis of individual differences in multidimensional scaling via an N-way generalization of “Eckart-Young” decomposition. Psychometrika 1970;35(3):283–319.

[5] Robert P, Escoufier Y. A unifying tool for linear multivariate statistical methods: the RV-coeffιcient. Journal of the Royal Statistical Society: Series C (Applied Statistics) 1976;25(3):257–265.

[6] Smilde AK, Kiers HA, Bijlsma S, Rubingh C, Van Erk M. Matrix correlations for high-dimensional data: the modified RV-coeffcient. Bioinformatics 2008;25(3):401–405.

[7] Foygel R, Drton M. Extended Bayesian information criteria for Gaussian graphical models. In: Advances in neural information processing systems; 2010. p. 604–612.

[8] de Rooi J, Nørgaard SK, Rasmussen MA, Bønnelykke K, Bisgaard H, Smilde AK. Data representations and-analyses of binary diary data in pursuit of stratifying children based on common childhood illnesses. PloS one 2018;13(11):e0207177.

[9] Bisgaard H, Vissing NH, Carson CG, Bischoff AL, Følsgaard NV, Kreiner-Møller E, et al. Deep phenotyping of the unselected COPSAC 2010 birth cohort study. Clinical & Experimental Allergy 2013;43(12):1384–;1394.

[10] Friedman J, Hastie T, Tibshirani R. Sparse inverse covariance estimation with the graphical lasso. Biostatistics 2008;9(3):432–;441.

[11] Mazumder R, Hastie T. The graphical lasso: New insights and alternatives. Electronic journal of statistics 2012; 6:2125.

[12] Su W, Bogdan M, Candes E, et al. False discoveries occur early on the lasso path. The Annals of statistics 2017;45(5):2133–;2150.

[13] Revai K, Mamidi D, Chonmaitree T. Association of nasopharyngeal bacterial colonization during upper respiratory tract infection and the development of acute otitis media. Clinical Infectious Diseases 2008;46(4):e34–;e37.

[14] Sevelsted A, Stokholm J, Bønnelykke K, Bisgaard H. Cesarean section and chronic immune disorders. Pediatrics 2015;135(1):e92–;e98.

[15] Lux AL, Henderson A, Pocock SJ, et al. Wheeze associated with prenatal tobacco smoke exposure: a prospective, longitudinal study. Archives of disease in childhood 2000;83(4):307–;312.

[16] Lim RH, Kobzik L, Dahl M. Risk for asthma in offspring of asthmatic mothers versus fathers: a meta-analysis. PloS one 2010;5(4).

[17] Litonjua AA, Carey VJ, Burge HA, Weiss ST, Gold DR. Parental history and the risk for childhood asthma: does mother confer more risk than father? American journal of respiratory and critical care medicine 1998;158(1):176–;181.

[18] Pedersen T. tidygraph: A Tidy API for Graph Manipulation. tidygraph: A Tidy API for Graph Manipulation 2018;.

[19] Pedersen T, ggraph: An Implementation of Grammar of Graphics for Graphs and Networks. 2018; 2018.

[20] Csardi G, Nepusz T, The igraph software package for complex network research. InterJ Complex Syst. 2006; 1695; 2018.

